# Histone 4 lysine 5/12 acetylation provides a plasticity code with epigenetic memory of environmental exposure

**DOI:** 10.1101/2022.07.21.500386

**Authors:** Michael S. Werner, Tobias Loschko, Thomas King, Tobias Theska, Mirita Franz-Wachtel, Boris Macek, Ralf J. Sommer

## Abstract

Development can be altered to match phenotypes with the environment, and the genetic mechanisms that direct such alternative phenotypes are beginning to be elucidated^1,2^. Yet, the rules that govern environmental sensitivity *vs*. invariant development (canalization), and potential epigenetic memory, remain unknown. Here, we show that plasticity of nematode mouth forms is determined by histone 4 lysine 5 and 12 acetylation (H4K5/12ac). Acetylation in early larval stages provides a permissive chromatin state at specific switch genes, which is susceptible to induction during the critical window of environmental sensitivity. As development proceeds deacetylation shuts off switch gene expression to end the critical period. We show that inhibiting deacetylase enzymes leads to long-term epigenetic memory, demonstrating that histone modifications in juveniles can carry environmental information to affect organismal traits in adults. This epigenetic regulation of plasticity appears to be derived from an ancient mechanism of licensing developmental speed that is conserved between flies and nematodes. Thus, H4K5/12ac provides a histone ‘plasticity’ code with epigenetic potential that can be stored and erased by acetylation and deacetylation, respectively.

**Highlights:**

1. Reciprocal transplant experiments reveal a critical time window of mouth-form plasticity.
2. Entry and exit of the critical window is determined by H4K5/12ac at the switch gene *eud-1*.
3. H4K12ac maintains transcriptional competence by supporting elongation.
4. Inhibition of deacetylation freezes an initial developmental trajectory, resulting in long-term epigenetic memory.
5. H4K5/12 acetylation control of plasticity was co-opted from an ancestral role in controlling developmental speed.

## Results

Different environments can elicit distinct phenotypes from a single genotype, referred to as phenotypic plasticity^3,4^. Ecological and theoretical approaches over the last 50 years have formalized the evolutionary implications and significance of plasticity^5-8^. More recently, molecular approaches are honing in on the mechanisms that direct environmental influence^1,9^. In contrast, the mechanisms that provide environmental sensitivity remain unknown. Here, we use the experimentally tractable mouth dimorphism in *Pristionchus* nematodes to uncover the molecular determinants of plasticity. Adult *P. pacificus* worms express either a narrow Stenostomatous (St) morph with a single dorsal ‘tooth’, or a wide Eurystomatous (Eu) morph with two teeth^2,10^. Morphological plasticity is coupled to behavioral plasticity, as adult St animals are strict bacterial feeders, while Eu animals can use their opposing teeth to kill other nematodes for food or competitive advantage^11-15^ (Fig. 1a,b). We found that specific histone modifications - acetylation of histone 4, lysines 5/12 - establishes a ‘plasticity code’ at switch genes that provides epigenetic memory of environmental experience. This code also regulates developmental rate across multiple Ecdysozoan species. Thus, these results reveal the molecular determinants of plasticity and their potential evolutionary origin.

**Fig. 1.**
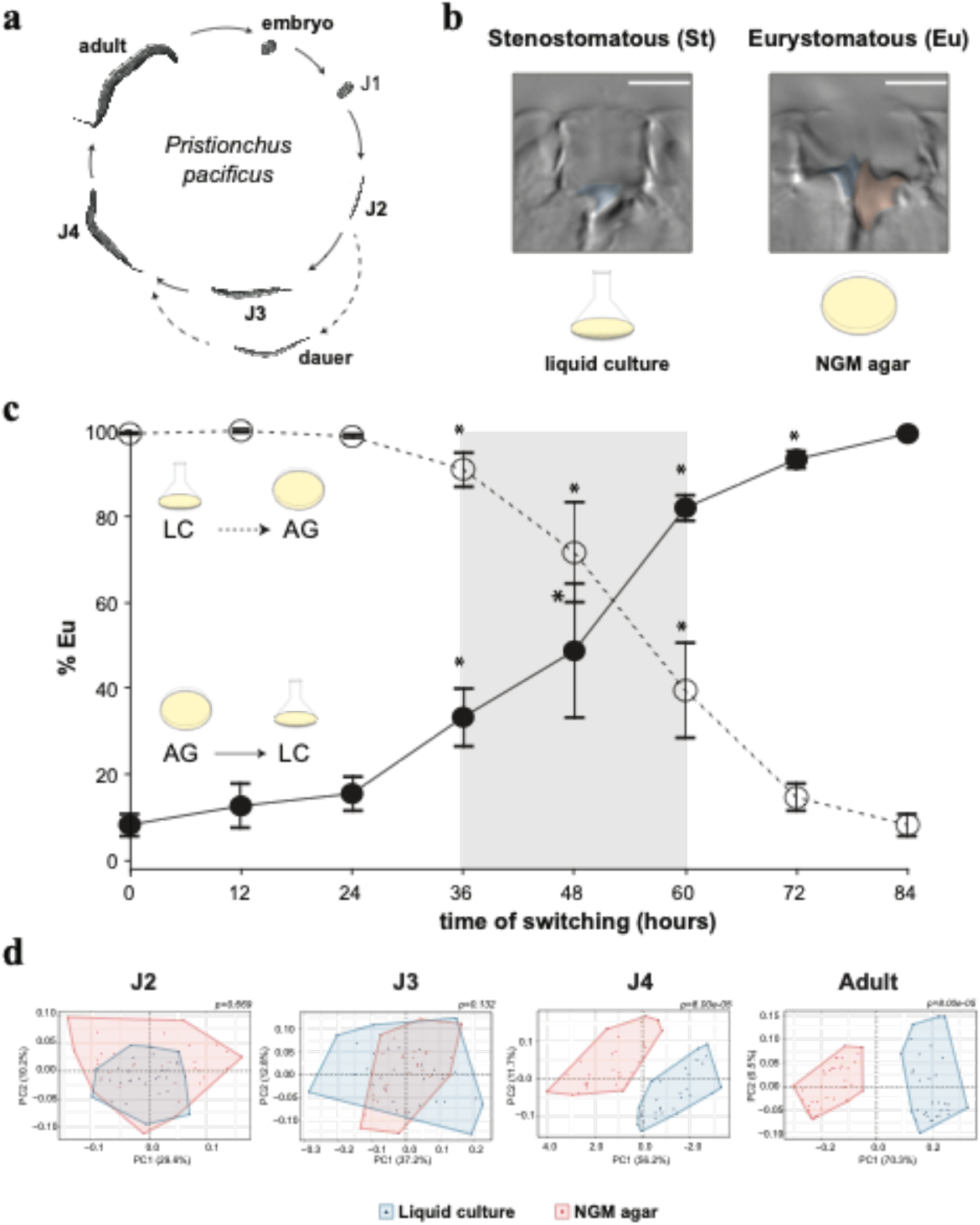
Mouth-form phenotypic plasticity is determined during a critical window in development. **a**, Life cycle of *Pristionchus pacificus*. **b**, Stenostomatous (St, left) and Eurystomatous (Eu, right). Scale bar = 5 µm. **c**, Switching experiments delineate developmental phases of plasticity. Adult phenotypes are plotted as a function of transferring between environments after synchronization. Statistical significance was assessed by binomial logistic regression on Eu and St counts between a given time-point, and the t’=0 and 84 hour phenotype; *=p<0.05 relative to both. Error bars represent S.E.M. for n=5 independent biological replicates. Note, intermediate values are intermeidate *ratios*, not intermediate morphs. **d**, GMM analysis of mouth-shape differences throughout development. PCA plots show the morphospace that contains shape variation per stage. Effect size (Z) for J2= 1.403, J3= 2.172652, J4= 3.847113, Adult= 5.0949. p-values represent FDR-adjusted pairwise Procrustes Anova.

### Reciprocal Transplantation Reveals a Critical Window of Environmental Influence

In the laboratory, different culture conditions can be harnessed to bias morph development (Fig. 1b): the wild type *P. pacificus* strain PS312 is predominantly St when grown in liquid S-Medium and Eu when grown on NGM agar plates^16^. We investigated the boundaries of environmental sensitivity by performing reciprocal transplant experiments between liquid and NGM-agar. Transferring between environments at 12 and 24 hours after egg synchronization (t’0) led to a complete adoption of the second environment’s phenotype, regardless of the direction of the environmental shift (Fig. 1c; n=5, *p*<0.05). Thus, this period represents a naïve and fully plastic phase of development. In contrast, when animals were switched at ≥ 36 hours they began to retain memory of their previous environment. Specifically, an increasing percentage of animals executed the morph of the first environment (Fig. 1c). Finally, after 60 hours, transplanting had little to no effect on mouth-form ratios. These results reveal a critical time window of environmental sensitivity between 36 and 60 hours. Prior to that, juveniles are completely plastic while afterwards the decision is irreversible.

### Molecular and morphological plasticity underly the critical window

Next, we looked into potential mechanisms that provide plasticity during the critical window and end plasticity after it. ‘ Switch genes’ determine alternative phenotypes by being expressed above or below a given threshold^17,18^. *eud-1* is a conserved steroid sulfatase that yields 100% Eu animals when constitutively overexpressed and 100% St animals when knocked out^19^. We predicted that transcriptional plasticity of switch genes (i.e., *eud-1)* underlies phenotypic plasticity during the critical window, while invariant expression demarcates developmental canalization. To test this prediction, we measured mRNA levels of *eud-1* during normal development and reciprocal transplant experiments. At 12 hours, we found strong induction of *eud-1* in NGM-agar (Eu condition) compared to liquid culture (St condition)(Extended Data Fig. 1a,b). Surprisingly though, modest expression was still observed in liquid culture, indicating some amount of environment-insensitive transcription during the naïve phase. This modest expression is rapidly induced when worms are transferred to NGM-agar during the critical window, or decreased when transferred from NGM-agar to liquid (Extended Data Fig. 1a,b; p<0.05), demonstrating environmental sensitivity. Intriguingly, *eud-1* begins to be repressed after 36 hours regardless of the environmental condition, such that at 60 hours *eud-1* mRNA levels are normalized between environments – coinciding with the end of the critical window. Thus, plasticity is connected to switch gene flexibility and exit from the critical window is connected to switch gene repression.

Next, we wondered if plasticity during the critical window is confined to a specific juvenile stage/molt. First, we measured juvenile development after hypochlorite-synchronization and compared them to our reciprocal transplant data, which indicated that plasticity peaks in the J3 stage (Extended data Fig. 1c-g). Second, fitting our reciprocal transplant data to a logistic model revealed inflection points between 48-54 hours, which coincides with the J3-J4 molt (Extended data Fig. 1h,i). Third, quantitative geometric morphometrics (GMM) throughout development revealed significant differences between conditions beginning in the J4 stage (Fig. 1d, p <0.05 and Z≥2.0; Extended Data Fig. 2), even though mouth dimorphism was previously thought to occur only in adults^20^. Collectively, these data demonstrate that switch gene transcription is indeed sensitive during the critical window, while *eud-1* repression and morphological differentiation correspond with the end of it - providing a plausible mechanism for the establishment of environmentally sensitive critical periods of development.

### Morph development is regulated by H4K5/12 acetylation

We hypothesized that open and closed chromatin structure – mediated by histone modifications^21-23^ - underlies transcriptional plasticity *vs*. repression of *eud-1*. First, we queried a panel of inhibitors that target histone-modifying enzymes for a potential effect on mouth form (Fig. 2a,b). Interestingly, the Tip-60/KAT5 histone acetyltransferase inhibitor NU9056 converted normally Eu animals on NGM-agar to the St morph (*p*=1e-15), while the histone deacetylase (HDAC) inhibitor Trichostatin A (TSA) converted normally St animals in liquid culture to the Eu morph *(p=4*.*3e-* 12). Hence, functionally opposite inhibitors yielded correspondingly opposite phenotypes, indicating that histone acetylation/deacetylation has an important role in mouth-form development.

**Fig. 2.**
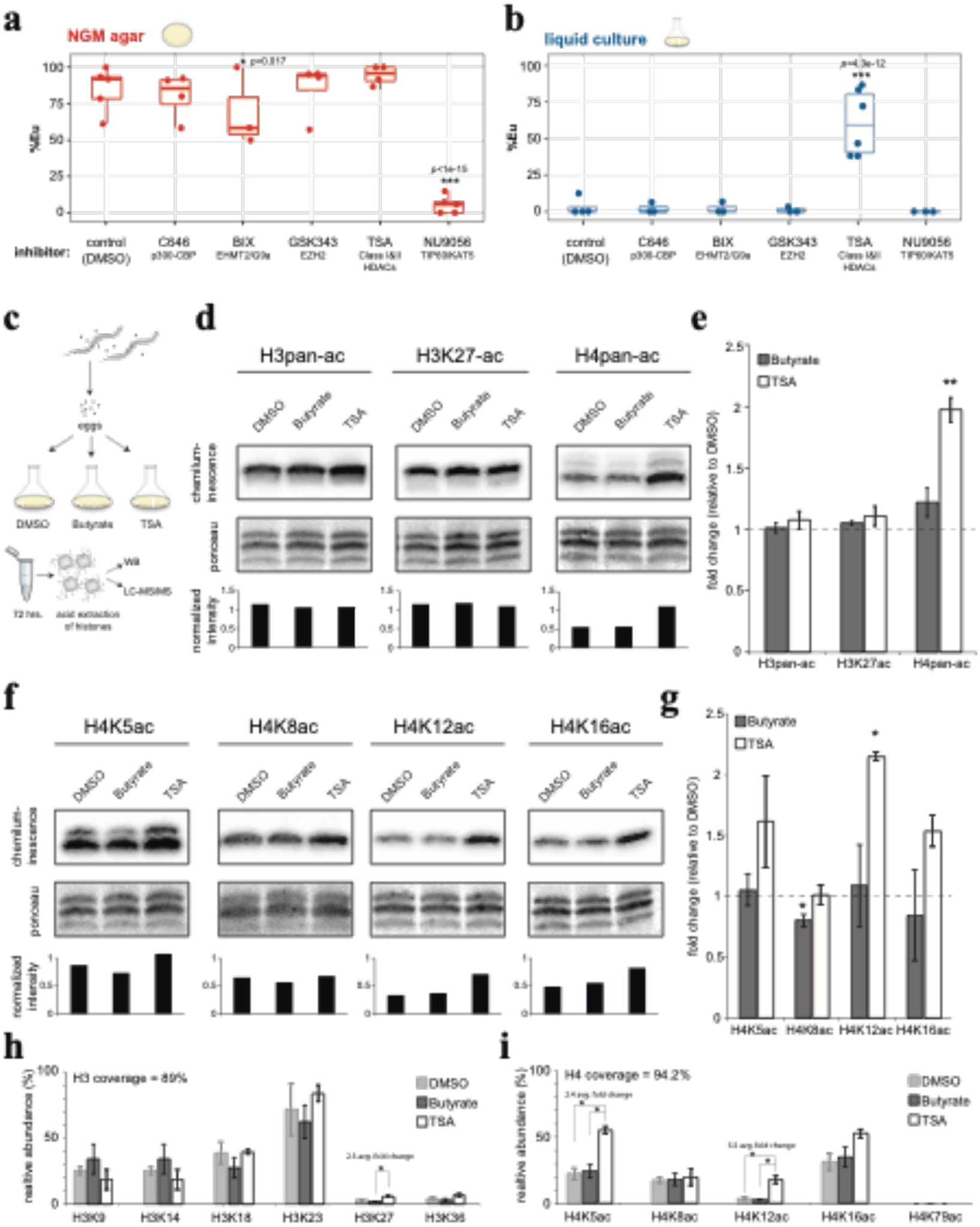
Mouth form is linked to H4K5/12ac. **a, b**, Phenotype (%Eu) after exposure to histone-modifying enzyme inhibitors in each environment. Target enzymes indicated below inhibitors. Box-plot upper and lower boundaries represent the 25% and 75% quantile, respectively, middle bar represents 50% quantile (median). Statistical significance determined by binomial logisitic regression on Eu and St counts. **c**, Strategy to identify the molecular mechanism of TSA’s effect: Bleach-synchronized eggs were seeded into liquid culture with DMSO (negative control for solvent), Butyrate (negative control for HDAC inhibitor that had no effect on phenotype), and TSA. Worm pellets were used for acid-extraction of histones and subject to WB and LC-MS/MS. **d**, Representative WBs of *P. pacificus* histones with specified antibodies, total histone staining by ponceau, and resulting relative intensities by quantitative densitometry. **e**, Mean fold change vs. DMSO, n=3. Error bars represent S.E.M. Statistical significance was assessed by a two-tailed student’s t-tests between TSA and Butyrate, *=p<0.05, **=p<0.01. **f**,**g**, Same as d,e but with H4-specific antibodies. **h**, Relative H3 and **i**, H4 acetylation of indicated residues compared to total H3 and H4 peptide intensities. Statistical signficance was assessed by a student’s t-test.

NU9056-treated animals had pleiotropic effects (e.g., reduced fertility, egg laying) consistent with *Ppa-mbd-2* and *Ppa-lys-12* mutants that are egg laying- or vulva defective and are St-biased^24^. However, animals that were exposed to TSA appeared wild type except for the effect on mouth form. HDAC inhibitors have also been shown to affect developmental decisions in other organisms, and several are currently in use as chemotherapeutic drugs^22,23^’^25^’^26^. Yet, it is not clear what the downstream targets and mechanisms of action are. Therefore, in the following we focus on TSA and the potential role of histone deacetylation in plasticity.

We confirmed that TSA also induced the Eu morph in axenic liquid culture, arguing for a direct inhibition of nematode enzymes rather than an indirect effect on their bacterial diet (p=1.2e-6)(Extended Data Fig. 3a). TSA also increased the proportion of Eu animals in three different *Pristionchus* species (Extended Data Fig. 3b), indicating that acetylation has an evolutionary conserved role in mouth form across the genus. Surprisingly however, although TSA is a pan-deacetylase inhibitor^27^, no obvious effect on mouth form was seen after treatment with several other HDAC inhibitors (Extended Data Fig. 3c). Presumably this is due to an unusual degree of specificity between TSA and its target enzyme in *Pristionchus*, and we wondered if that would allow us to investigate the role of discrete acetylated residues in plasticity.

To determine whether specific H3 and H4 residues are affected by TSA in *P. pacificus* we performed Western Blots (WB) on acid-extracted histones (Fig. 2c, Extended Data Fig. 4). In addition to DMSO (solvent) we used butyrate as a second negative control, because it is an HDAC inhibitor that did not affect mouth form (Extended Data Fig. 3c). H3K27ac is a reproducible marker of active enhancers and promoters^21^, and has recently been implicated in behavioral differences between ant castes^22,28^. However, we did not observe an increase in H3 acetylation using a pan-acetyl antibody or with a specific antibody toward H3K27ac (n=3, Fig. 2d,e). In contrast, we observed a 2-fold induction of H4 acetylation (±0.1, p=6.3e-4 *vs*. DMSO and 8.5e-3 *vs*. butyrate). To verify TSA’s effect on H4 and to determine which H4 lysine(s) are hyperacetylated, we repeated WBs with specific H4-acetyl antibodies (Fig. 2f,g). Consistent with the apparent specificity of TSA in *Pristionchus*, we observed only one H4 lysine with significant hyperacetylation relative to both controls: H4K12 (p=5.7e-6, 3.6e-2, respectively).

To further validate this result, we performed Liquid Chromatography Tandem-Mass Spectrometry (LC-MS/MS) (n=2, Fig. 2c, and h,i, Extended Data Table 2). In agreement with our WB data, H4K12ac exhibited the greatest increase in TSA for all queried H3 and H4 lysine residues (avg. fold change = 5.5). The greater sensitivity of LC-MS/MS revealed a smaller yet significant increase in H4K5ac against both DMSO and butyrate (avg. fold change = 2.4). No other modification was statistically significant against both controls (e.g., H3K27ac exhibited an increase relative to butyrate, but not to DMSO). Collectively, both immunostaining and mass spectrometry implicate H4 acetylation, and in particular H4K12ac, as the main effector by which TSA influences mouth-form development.

### An H4 acetylation/deacetylation timer determines the critical window

The role of H4K12ac is poorly understood, but there are intriguing links to aging^29^, hormone-dependent cancer^30^, and learning^31^; processes that are all connect to plastic gene regulation. Chromatin Immunoprecipitation (ChIP) experiments in human cells showed that H4K12ac is distributed throughout gene bodies in comparison to other acetylated lysines that demarcate enhancers or promoters^21,32^, hinting at a role in transcription elongation rather than initiation. Using RT-qPCR, we found that TSA increased *eud-1* expression compared to DMSO (Fig. 3a). However, instead of inducing *eud-1*, TSA appeared to prevent the repression that normally occurs after 36 hours. These data suggest that H4K5/12ac does not have a role in environmental induction, but rather maintains transcriptional competence in the critical window. Then, at 36-48 hours H4K5/12 deacetylation leads to switch gene repression. These findings differentiate H4K5/12ac from other histone acetylation marks that are thought to directly promote transcriptional initiation^33^.

**Fig. 3.**
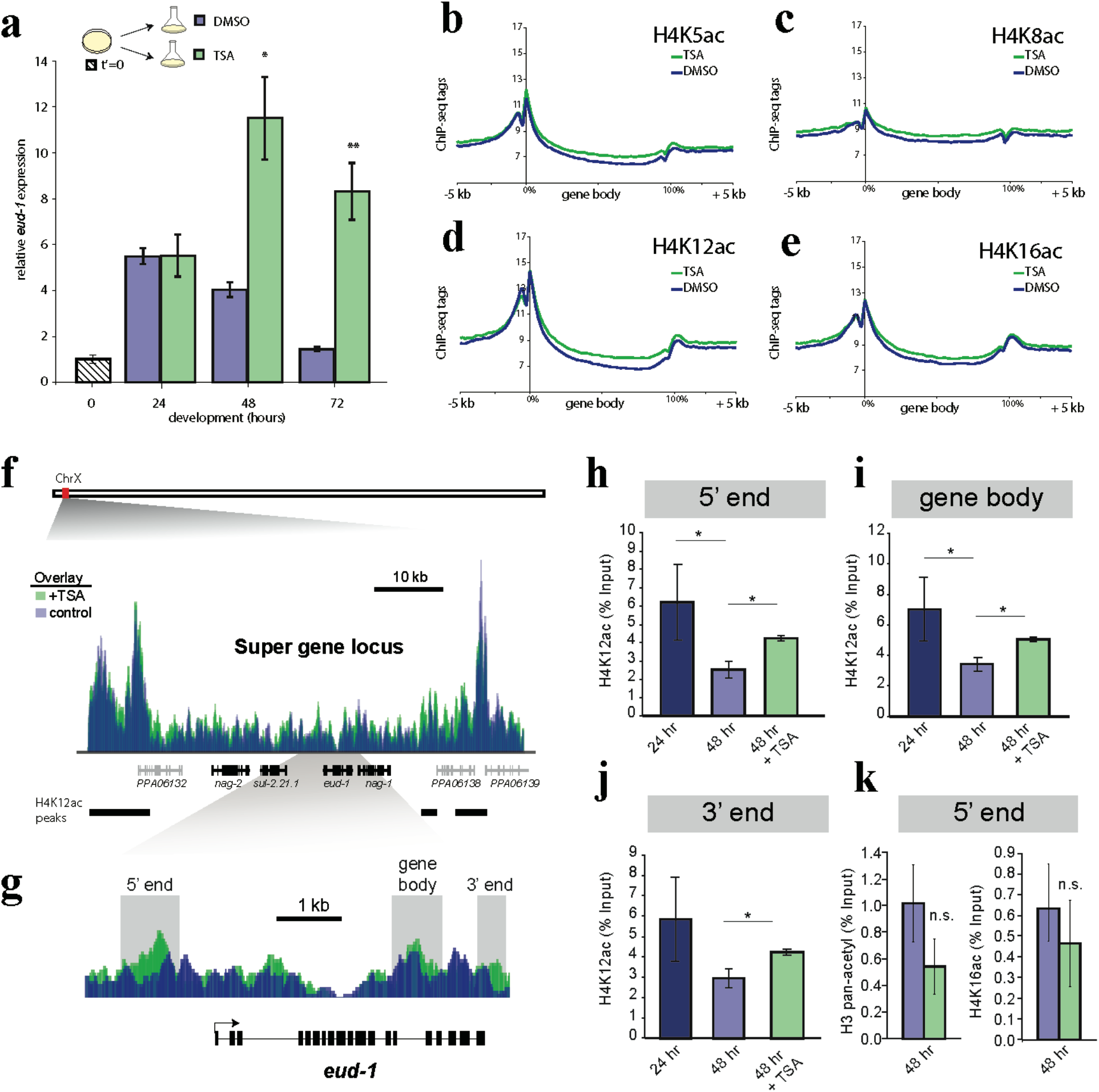
H4K12ac acetylation/deacetylation at switch genes determines plasticity and canalizaiton, respectively. **a**, *eud-1* expression at indicated time points by RT-qPCR with either DMSO or TSA added directly after bleach synchronization. Error bars represent S.E.M. of 3-4 biological replicates. Statistical significance was determined by student’s t-test, *=p<0.05, **=p<0.01. **b-e**, ChIP-seq meta-gene profiles of H4-acetylated residues at 48 hours in +/-TSA. y-axis represents average input-normalized ChIP-seq tag counts per bp per gene, and the x-axis represents 5’ to 3’ gene coordinates divided into 200 bins +/-5 kb, n=2. **f**, ChIP-seq tag density of H4K12ac over the multi-gene locus (black genes) that control mouthform plasticity, which are flanked by large peaks. y-axis = 0-72 depth-normalized density for both tracks. Black bands indicate H4K12ac peaks in liquid culture determined by MACS2. **g**, Zoom in on *eud-1* and regions targted by qPCR in h-k. **h-j**, ChIP-qPCR of H4K12ac across *eud-1* at 24 hours (dark blue), 48 hours (light blue) and 48 hours when grown in TSA (green), n=3. **k**, ChIP-qPCR of H3-pan-acetyl and H4K16ac at the 5’end of *eud-1*, n=3.

To explore this possibility further, and to examine comprehensive patterns of H4 acetylation, we performed ChIP-seq with antibodies targeting H4K5ac, H4K8ac, H4K12ac, and H4K16ac at 48 hours in +/-TSA conditions. Consistent with our original biochemical results (Fig. 2), H4K12ac exhibited the greatest induction in TSA, followed by H4K5ac (Fig. 3b-e). Subtle increases could also be seen with the other H4ac modifications, although it is formally possible that this is a result from antibody cross-reactivity^34^. Interestingly, the increase in H4K12ac was almost exclusively over gene bodies rather than at promoters, which supports the hypothesized role in elongation. This pattern also appeared to be true at the locus encompassing the *eud-1* switch gene, which is part of a four-member ‘super-gene’ on the X chromosome where each gene contributes to mouth form regulation^35^. Our ChIP-seq data revealed that the super-gene locus is bordered by large peaks of H4 acetylation (Fig. 3f, Extended Data Table 3), and that TSA led to an increase in H4K12ac between these peaks, including across *eud-1*, rather than at the peaks themselves (Fig. 3g).

Next, we performed ChIP-qPCR at 24 and 48 hours to see if we could track deacetylation across *eud-1* during the critical window. Indeed, we found a significant decrease in H4K12ac at 3/3 qPCR primer locations spanning the *eud-1* gene locus (Fig. 3h-j, p<0.05). However, adding TSA partially maintained the early, high levels of H4K12ac at 48 hours (Fig. 3h-j); arguing that TSA prevents *eud-1* repression by preventing H4 deacetylation. In contrast, we did not detect an increase in either H3 acetylation or H4K16ac at *eud-1* in the presence of TSA, confirming the specificity of our previous results (Fig. 3k). Collectively, these data lead toward a model of early H4K5/12ac deposition that primes transcription of switch gene(s) for environmental induction. During the J3-J4 transition, an unknown developmental mechanism leads to deacetylation, turning off switch gene expression and effectively closing the critical window (Extended Data Fig. 5).

### Epigenetic memory can be enforced by HDAC inhibition

The role of histone modifications in plasticity opens up the potential for epigenetic gene regulation; a long-held but poorly supported hypothesis to connect plasticity to adaptation^7^. To determine whether histone acetylation can provide epigenetic memory, we assessed if preventing deacetylation during the critical window would ‘fix’ or ‘freeze’ an initial developmental trajectory despite shifting to a different environment. In principle, this would provide compelling evidence that histone modifications can carry long-term environmental information to affect future developmental decisions. To test this premise, we combined transplant experiments with TSA treatment. Specifically, we transferred worms between NGM-agar and liquid at 24 hours, which normally leads to the St phenotype (Fig. 1c), but this time added TSA at the time of switching. Remarkably, these worms were phenotypically similar to having experienced the agar environment for their entire development (90.8% Eu ±1.2)(Fig. 4). Not only were environmentally shifted TSA-treated worms significantly different from DMSO controls, they were also significantly different from worms treated with TSA at t’=0 but without switching (60.9% Eu ±8.0). These results demonstrate a combined effect of the environment with HDAC inhibition that is consistent with fixing the initial Eu developmental trajectory. We then performed a series of experiments to attempt to refute this interpretation (Fig. 4). First, to rule out that simply adding TSA at a later time point has a greater effect on mouth form than at t’=0, perhaps due to degradation, we added TSA at 24 or 48 hours without environmental transfer. These experiments resulted in decreasingly intermediate ratios of Eu animals similar to, and less than adding TSA at t’=0, respectively (p<0.05). We also repeated transplant experiment at 24 hours, but waited until 48 hours to add TSA. Again, this experiment failed to induce a typical NGM-agar phenotypic ratio (28.5% Eu ±3.8). These results show that there is an additive effect of the environment with TSA, and that this effect depends on adding TSA at the precise time of environmental transfer. We conclude that preventing histone deacetylation after a temporary juvenile experience in NGM-agar fixed the Eu developmental trajectory; 48 hours and two molts prior to adult differentiation. Thus, epigenetic memory of environmental exposure can be stored and erased by acetylation and deacetylation, respectively.

**Fig. 4.**
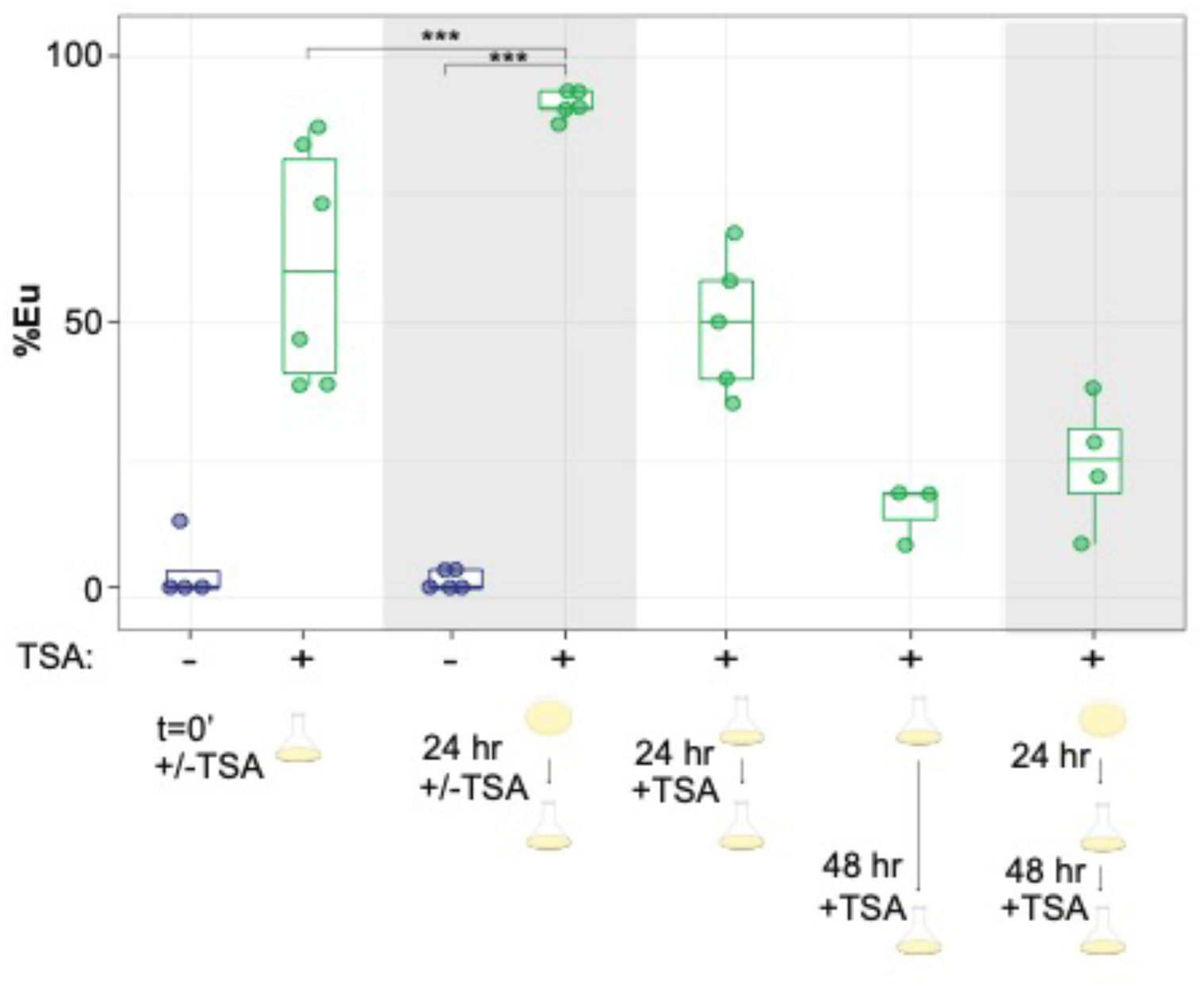
H4 acetylation can provide long term epigenetic memory. Transplant experiments between NGM-agar and liquid culture +/-TSA at different time points. When TSA is not added (-), the solvent DMSO was added. Individual data points are shown (n=3-6 biological replicates), and box-plot upper and lower boundaries represent the 25% and 75% quantile, respectively, and the middle bar represents the 50% quantile (median). Statistical significance determined by binomial logisitic regression on Eu and St counts, *=p<0.05, **=p<0.01, ***=p<0.001.

### TSA treatment delays development in worms and flies

The extended *eud-1* expression in TSA was reminiscent of repeated or delayed developmental programs seen in some heterochronic gene mutants^36^. It was also previously shown that the Eu morph develops ∼6 hours slower than the St morph^37^. Therefore, we speculated that acetylation/deacetylation might affect plasticity by affecting developmental speed. We coupled hypochlorite-treatment with temporary starvation to obtain highly synchronized cultures, and compared developmental rates in the presence or absence of HDAC inhibition. Strikingly, we found that TSA prolonged development by several hours (Fig. 5a). To explore the generality of this phenomenon, we repeated these experiments in *Caenorhabditis elegans*, and again found a similar developmental delay (Fig. 5b). Importantly, *C. elegans* lacks mouth-form plasticity, implying that the effect on development predates the effect on mouth form. Recent data suggest that the divergence time between *C. elegans* and *P. pacificus* is ∼200 million years ago^38^, hinting at a deeply conserved mechanism to control development timing. Indeed, when searching the literature, we found a similar effect of TSA on development had been reported in *Drosophila melanogaster* in 2003^39^, which we independently confirmed (Fig. 5c). Thus, HDAC inhibition delays development in three highly diverged species of Ecdysozoa, a superphylum of molting animals spanning hundreds of millions of years of evolution that potentially predates the Cambrian explosion^38^.

**Fig. 5.**
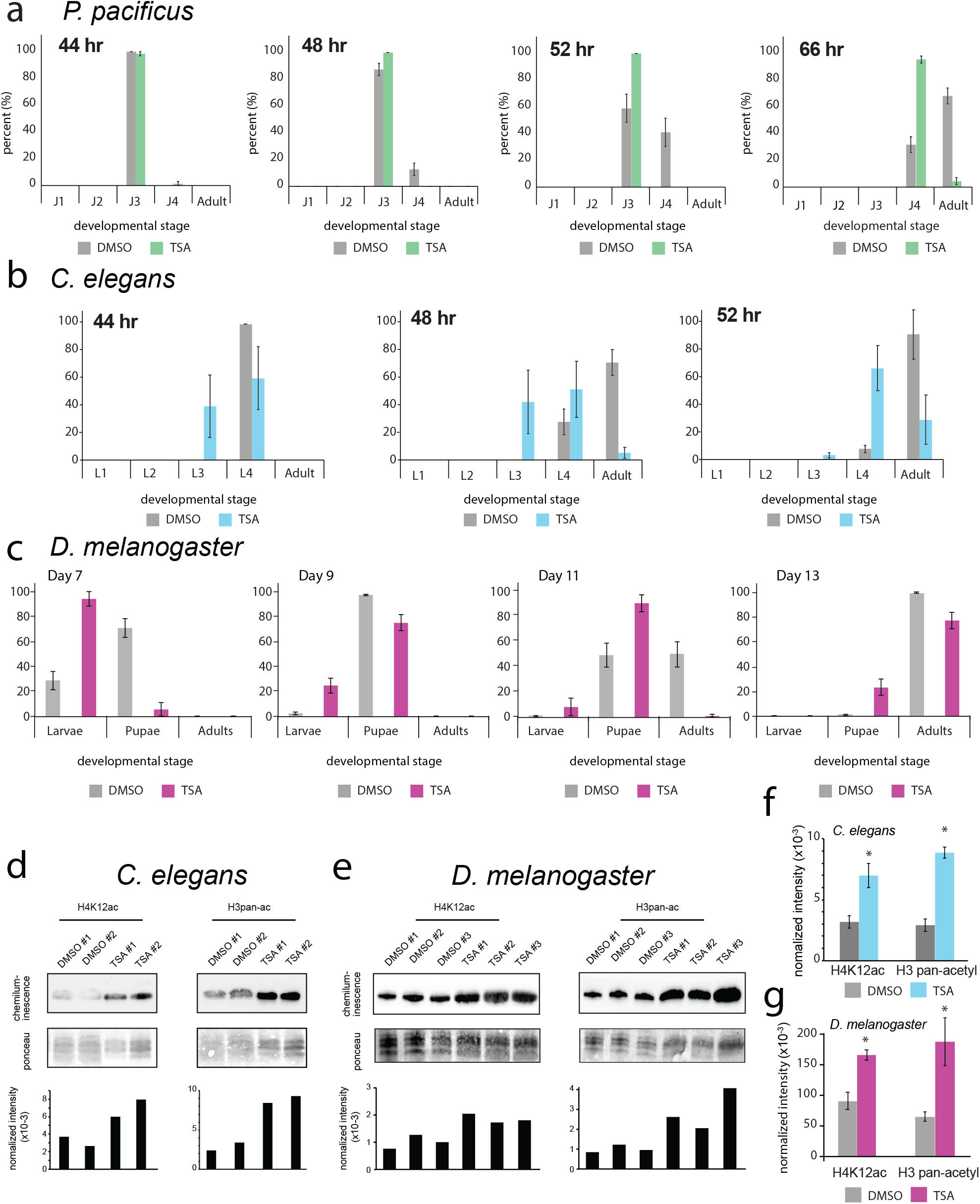
Histone acetylation links plasticity to developmental speed. **a**, Developmental rate of *P. pacificus*, **b**, *C. elegans*, and **c**, *D. melanogaster* +/-TSA, n =2 for nematodes and 4 for flies. **d**, Representative WBs of acid-extracted histones from *C. elegans* and **e**, *D. melanogaster*. Intensity of chemiluminescence was normalized to total histone levels detectedy by Ponceau S. **f**, Mean normalized intensity (n=3) of WBs in *C. elegans* and **g**, *D. melanogaster*. Statistical significance determined by 1-tailed students t-test.

### Histone acetylation may regulate plasticity by licensing developmental speed

Finally, we examined whether H4 hyperacetylation is the cause of delayed development. We found a 2.2-fold induction of H4K12ac in the presence of TSA in *C. elegans*, and a 2.6-fold induction in *D. melanogaster* (Fig. 5d-g, p<0.05), in good agreement with our results from *P. pacificus*. While H4K12 hyperacetylation is common to all species, we also saw an increase in H3 acetylation in *C. elegans* and *D. melanogaster*, consistent with previous reports that TSA can have pan-deacetylase activity^27^. Given that both mouth form and developmental timing in *P. pacificus* are regulated by deacetylation, but control of timing is ancestral, we hypothesize that developmental licensing by H4K5/12ac was co-opted for the evolution of mouth-form plasticity.

## Discussion

Phenotypic plasticity allows organisms to adjust development to match their environment^7^. Environmental sensitivity is limited to a specific stage of development, referred to as the critical ‘window’ or ‘period’. Here, we identify that entry and exit of the critical window is defined by a histone code of H4K5/12 acetylation and de-acetylation, and that H4 acetylation can provide epigenetic memory of previous experiences. Furthermore, we provide evidence that this axis is an ancient timer for regulating development speed.

The role of H4K5/12ac in plasticity was uncovered in part due to the unusual specificity of TSA in *P. pacificus*. H4 N-terminal tail acetylation is broadly correlated with gene activation^40^, however, the functions of most individual acetyllysines are not well known. H4K16ac is the exception, and has been shown to be necessary for dosage compensation in flies^40^ and hematopoietic differentiation in mammals^41^. However, we did not observe a significant correlation of this mark with switch gene transcription, or with commonly studied enhancer/promoter modifications on H3^21^. There is indirect evidence though for a role of H4K12ac in plasticity. For instance, H4K12ac has been linked to acetyl-CoA levels^29^, suggesting that it may connect diet or metabolism to changes in gene expression. Furthermore, stimulating H4K12ac can promote memory formation, a paradigm of neuronal plasticity^31,42^. While the outcomes of neuronal and morphological plasticity are different, the proximate mechanisms regulating plasticity may be shared.

Our time-resolved data also support a mechanistic role for H4K5/12ac in transcriptional elongation rather than initiation. This has been hinted at by ChIP-seq data showing an abundance of H4K12ac in gene bodies compared to other acetylated lysines, which show a bias toward promoters and enhancers^32,43^. Furthermore, the bromodomain and extraterminal domain (BET) family protein BRD4, which binds to acetylated histones, was recently shown to act as an elongation factor to facilitate RNAPII clearance through chromatin^44^. The related protein BRD2 interacts specifically with H4K5/12ac in immortalized human cell lines^45,46^, while BRD4 has been shown to promote estrogen receptor-positive breast cancer by interacting with hyperacetylated H4K12^30^. It will be fascinating to see if phenotypic plasticity follows similar processes, and if H4K12ac underlies TSA’s particular potency against breast cancer^47,48^. Alternatively, TSA can induce cell cycle arrest through transcriptional regulation of p21^49^, which may explain the delayed development we observed at the whole organism level. Untangling these mechanistic possibilities may provide insights into both fundamental developmental biology and drug design.

Histone modifications have been proposed to carry long-term environmental information. However, the half-life of histone acetylation is generally considered too short (1-2 hours) to provide epigenetic memory^50^. Nevertheless, more recent studies have shown that histone acetylation can persist through mitotic division^45,51^’^52^; reviving the question of their epigenetic potential. Our results add an important next layer by demonstrating that histone acetylation can provide epigenetic memory in a multicellular organism. Our results also provide empirical and mechanistic evidence that phenotypic plasticity is channeled through epigenetic regulation^7^.

Lastly, the confluence of developmental timing and plasticity has implications beyond gene regulation. Regulating developmental timing, referred to as heterochrony, has been hypothesized for nearly a century to facilitate evolutionary novelty^53^. While ample morphological evidence supports this hypothesis, there are scant known molecular mechanisms. Our results suggest that acetylation/de-acetylation acts as a timer to progress through development, which was utilized by *Pristionchus* to acquire the Eu morph by extending switch gene *(eud-1)* expression. If sufficiently selected upon, the Eu morph would ultimately canalize, as appears to have occurred in some *Pristionchus* species^54^, representing plasticity-first evolution^55^. Going forward, it will be important to evaluate how H4K5/12 acetylation affects both molecular and evolutionary mechanisms.

## Methods

### Environmental Switching Experiments

Gravid adult *P. pacificus* were bleach-synchronized ^56^ and eggs were seeded into agar or liquid media as described in Werner et al., 2017, and then switched to the other condition at the indicated time points. At 84 hours post-synchronization, adult worms were phenotyped with a 100x/1.4 oil DIC objective (Zeiss), and %Eu was plotted on the y-axis. Statistical significance was assessed by binomial logistic regression on Eu and St counts between a given time point and both the t’0 and the 84 hour’s phenotype (see section below on ‘Statistics on mouth-form frequencies’ for more details). Error bars represent S.E.M. for n=5 independent biological replicates.

### Statistics on mouth-form frequencies

To test the statistical significance of the effect of small molecule inhibitors or switching experiments on mouth-form frequencies, while incorporating biological replicates, we performed a weighted binomial logistic regression in the R base statistical software package. Eu counts were modelled as ‘successes’ and St modelled as ‘failures,’ and replicates were included as repeated measurements:

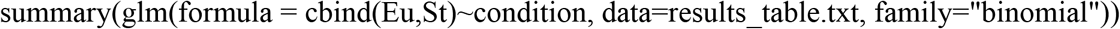

Resulting *p* values were adjusted for multiple testing using a *bonferroni-correction* (i.e., by the number of inhibitors tested), and are indicated in corresponding figure panels. *n* ≥ 3 in all cases, with individual replicates shown as data points.

### Fitting reciprocal transplant data

Raw data of transplantation experiments were normalized on the y-axes to ‘% plasticity’ by subtracting the average minimum value (e.g., the average %Eu when grown for the entire duration of development in liquid) from each time point. The normalized data were fit to a four-parameter logistic function, which is commonly used in pharmacology to fit a physiological response to drug concentration^57^ or ligand binding to a protein in biochemistry (i.e., the Hill equation):

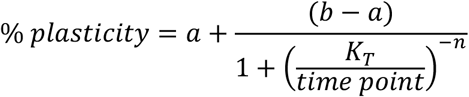

Where *a* = the minimum asymptote (or plasticity after determination), *b* = maximum asymptote (or plasticity at t’=0), *K*_*T*_ = the inflection point of the curve (or the time point when plasticity transitions to determination, equivalent to the effective concentration *EC*_*50*_ in pharmacological studies, or *K*_*A*_ in protein-binding experiments), and *n* = the slope at the steepest part of the curve (or the rate of determination, also known as the Hill coefficient). Note, *K*_*T*_ is not necessarily equivalent to 50% plasticity because the maximum asymptote or maximum plasticity *b* is not necessarily equal to 100%. To fit the data, a nonlinear least squares regression was performed in R using the ‘nls’ function with upper bound=100, lower bound=0, and starting estimates of *a* = 0, *b* = 100, *K*_*T*_ = 60, and *n* = 10, which reached convergence after four iterations. The fitted values were plotted using ‘ggplot2,’ and a line marking the inflection point *K*_*T*_ was added by ‘geom_vline.’

### Geometric morphometric (GMM) quantification

Two-dimensional mouth shapes were quantified using 13 fixed landmarks and 3 sliding semilandmarks. Landmark configurations from n ≥ 30 animals per stage and condition were used as input for shape analysis by General Procrustes Alignment (GPA), and PCA was performed on the aligned Procrustes coordinates (i.e., shapes) as described in Theska et al., 2020^58^. GPA was performed using ‘gpagen’ of geomorph (ver. 4.0.0); sliding of the semilandmarks was achieved by minimizing bending energy. Statistical testing for differences between group means of shape variation was performed on Procrustes coordinates of animals of the same stage in different conditions via permutational Multivariate Analysis of Variance *(PERMANOVA, or* ‘Procrustes’ ANOVA) with a randomized residual permutation procedure (RRPP). This was done using the ‘procD.lm’ function of geomorph with 100,000 iterations, seed = NULL. Resulting relative effect sizes (*Z*-scores) and FDR-adjusted *p*-values are presented adjacent to corresponding PCA plots. We considered a PERMANOVA effect as ‘incompatible with the null hypothesis’ if the relative effect size was greater than or equal to two times the standard deviation (i.e., *Z* ≥ 2.0) and the associated *p*-value was below a type I error rate of 0.05 (i.e., *p* < 0.05).

### Histone-modifying-enzyme inhibitor assays

Worm cultures were synchronized by adding bleach solution (1 ml bleach/0.5 ml 5M NaOH) to 3.5 ml of gravid adult worms, which were washed with M9 from 3 × 6 cm NGM agar plates per experimental condition. Eggs were then aliquoted to 10 ml S-medium liquid cultures or 6 cm NGM agar plates. For liquid experiments, inhibitors were added directly at the time of aliquoting eggs. For agar experiments, fresh 6 cm NGM plates were spotted with 500 µl of overnight cultures of *E. coli* OP50 (37°C in LB, 180 rpm) plus inhibitors and spread evenly over the surface of the plate, then air-dried in a chemical hood before adding 100-500 bleached eggs. The following concentrations of inhibitors or control (DMSO) were added: 100 µl DMSO (Sigma Aldrich cat. #D8418) = 1% (v/v), 100 µl 6.6 mM TSA (Selleckchem cat. #S1045) dissolved in DMSO = 66 µM final conc., 100 µl 2.2 mM C646 (Sigma Aldrich cat. # 382113) dissolved in DMSO = 22 µM final conc., 100 µl 1.7 mM BIX (Sigma Aldrich cat. #B9311) dissolved in DMSO = 17 µM final conc., 100 µl 1.85 mM GSK343 (Sigma Aldrich cat. #SML0766) dissolved in DMSO = 18.5 µM final conc., and 30 µl 43 mM NU9056 (Tocris cat. #4903) dissolved in DMSO = 129 µM final conc.. For Fig. S2, inhibitors were resuspended at the following concentrations before titrating into LC: 1 M Na-butyrate (Sigma Aldrich cat. #B5887) dissolved in water, 1 mM apicidin (Sigma Aldrich cat. #A8851) dissolved in DMSO, 10 mM CI-944 (Sigma Aldrich cat. #C0621) dissolved in DMSO, 10 mM TMP195 (Selleckchem cat. #S8502) dissolved in DMSO, 10 mM Pyroxamide (Sigma Aldrich cat. #SML0296) dissolved in DMSO, 5 mM SAHA/Vorinostat (Sigma Aldrich cat. #SML0061) dissolved in DMSO, 4 mM Tubacin (Sigma Aldrich cat. #SML0065) dissolved in DMSO, 10 mM CUDC-101 (Selleckchem cat. #S1194) dissolved in DMSO, 10 mM Panobinostat (MedChemExpress cat. #HY-10224) dissolved in DMSO, and 10 mM Belinostat/PXD101 (MedChemExpress cat. #HY-10225) dissolved in DMSO.

### Histone Purification and Western Blot

For nematodes, crude nuclei were obtained as in Werner et al. 2018^59^ but without sucrose cushion purification, with starting inputs of 200-500 µl worm pellets (10-20 × 10 cm plates of bleach-synchronized worms) collected at 72 hours in the presence of 100 µl DMSO, 66 µM TSA, or 10 mM butyrate (see ‘Histone-modifying-enzyme inhibitor assays’ method section for resuspension and cat. #s). Histones were acid-extracted from crude nuclei, and precipitated in TCA (Sigma Aldrich cat. #T9159) following Shechter et al., 2007^60^, and re-suspended in 80 µl water. Histone yields from each purification were determined by using a calibration curve of recombinant H3 and H4 (NEB, cat. #M2503S, #M2504S, respectively). Absolute amounts of histone were determined by densitometry on coomassie-stained bands with Fiji^61^ for *Pristionchus* data (conducted at the MPI in Tübingen) and Image Lab software (BioRad) for *C. elegans* and *D. melanogaster* (conducted at the University of Utah). For Western Blot, 5 µg of histone sample was loaded per lane on a BioRad ‘any Kd’ Precast gel (cat. #4569033) alongside 5 µl PageRuler Prestained Protein Ladder (Thermo Scientific cat. #26616), and run 200V for 28 minutes. A Wet Transfer was performed with Bjerrum Schaffer-Nielsen Buffer + SDS (48 mM Tris, 39 mM glycine, 20% methanol, 0.0375% SDS, pH 9.2) in a Mini Protean Tetra box (BioRad) to 0.2 µM, 7×8.5 cm precut nitrocellulose membranes (BioRad, cat. #162-0146), at 4°C with magnetic stirring and an opposing cold-pack, 100V for 10 minutes, then 60V for 20 minutes. Total-protein transferred was visualized by 10-20 ml 0.1% ponceau/5% acetic acid for 5-10 minutes with rotation, washed with distilled water until bands became apparent, and then imaged on a Quantum gel imager (Vilber Lourmat) with white light using the ‘preview’ function (in Tübingen) or BioRad ChemiDoc (Utah). Nitrocellulose blots were then briefly washed in TBS (50 mM Tris-HCl, 150 mM NaCl, pH 7.5), then blocked with 20 ml 5% nonfat dry milk (BioRad, cat # 170-6404) in TBS for 1 hour with rotation. Membranes were then washed 2 × 5-10 minutes in TBS plus 0.05% Tween 20 (TBS-T). Primary antibodies (Extended Data Table 1) were then incubated at 1:1,000-2,000 dilution in 5 ml 2.5% nonfat dry milk/TBS-T with membranes overnight (∼12 hours) at 4°C with rotation. The next day, membranes were washed 4 x with TBS-T, 5 minutes each with rotation. Secondary antibodies corresponding to the animal immunoglobulin of the primary antibody, fused to horseradish peroxidase (Anti-rabbit IgG-HRP, Cell Signalling, cat. #7074S), were then incubated at 1:2,000 in 5 ml 2.5% nonfat dry milk/TBS-T for 1 hour at room temperature. Membranes were then washed 4 x with TBS-T, plus one additional wash in TBS to remove residual Tween 20. To image, membranes were incubated in a 5 ml 1:1 A:B solution of Clarity Western ECL substrate (BioRad cat. #170-5060) for 5 minutes at room temperature with rotation, and chemiluminescence was detected on a Fusion SL imager (Vilber Lourmat; Tübingen) or ChemiDoc (BioRad; Utah) within 5 minutes.

For *Drosophila* experiments, flies were cultured on standard medium containing cornmeal, yeast, agar, and molasses, and maintained at 25°C and 60% humidity on a 16:8 light:dark cycle. Additionally, the medium contained either TSA dissolved in DMSO, or DMSO only as a control. These were added to the medium at a concentration of 0.34% (v/v) once it had cooled to approximately 50°C. This resulted in a final concentration of 10 µM TSA in the TSA-containing medium. The genotype of all flies used was *w*^*1118*^ (Bloomington Drosophila Stock Center #3605). For crude histone purification, between 100 and 150 wandering third-instar larvae were collected per replicate, and three replicates per treatment were used. Larvae were rinsed twice in PBS, and histone extraction, purification, and quantification were performed as described above. Western blots were performed as described above, except only 1 µg of histone sample was loaded per lane, and images were acquired using a ChemiDoc MP Imaging System (BioRad). Image and statistical analysis were performed as described above, except two-tailed F tests were conducted to ensure that variances did not differ significantly between treatments, and one-tailed Student’s *t*-tests were used since evidence from nematodes indicated that TSA was expected to cause increased histone acetylation. Antibodies used: H3pan = Active Motif #39140, lot #34519009; H4pan = Active Motif #39926, lot #18619005; H4K12 = Millipore #04-119-S, lot #3766681. Intensities of chemiluminescent bands were quantified and normalized to intensities of histone bands from Ponceau staining. Statistical testing on normalized intensities was performed by a two-tailed *student’s t*-test in nematodes and one-tailed test in *Drosophila, n* = 3 independent biological replicates for all experiments.

### Histone Liquid Chromatography Tandem-Mass Spectrometry (LC-MS/MS)

Histones were extracted from 72-hour worm pellets as described for Western Blots (see above) for two independent biological replicates each of 1% DMSO, 10 mM Na-Butyrate, and 66 µM TSA. 50-100 µg of soluble histones were then reduced with 1 mM DTT/50 mM ammonium bicarbonate for 1 hour at room temperature, and then alkylated for 1 hour in the dark with 10 mM chloroacetamide/50 mM ammonium bicarbonate. The reaction was neutralized with 10 mM DTT for 30 minutes. A 10x Arg-C digestion buffer was then added to histones (1x=5 mM CaCl2, 0.2 mM EDTA, 5 mM DTT), which were digested with 1:50 Arg-C protease (Promega, cat. #V1881): histone protein at 37°C for 12-16 hours. Digest completion was assessed by running an analytical SDS-PAGE of digested sample with undigested control sample. When complete, the reaction was stopped by adding 10% trifluoroacetic acid to a final concentration of 0.5%. Digested histones were then purified on homemade desalting C18 stage-tips ^62^, and run on an Easy-nLC 1200 system coupled to a QExactive HF-X mass spectrometer (both Thermo Fisher Scientific) in three technical replicates per biological sample as described elsewhere ^63^ with slight modifications: peptides were separated with a 127-minute segmented gradient from to 10-33-50-90% of HPLC solvent B (80% acetonitrile in 0.1% formic acid) in HPLC solvent A (0.1% formic acid) at a flow rate of 200 nl/min. The 7 most intense precursor ions were sequentially fragmented in each scan cycle using higher energy collisional dissociation (HCD) fragmentation. In all measurements, sequenced precursor masses were excluded from further selection for 30 s. The target values for MS/MS fragmentation were 10_5_ charges and 3×10^6^ charges for the MS scan.

Data was analyzed by MaxQuant software version 1.5.2.8^64^ with integrated Andromeda search engine^65^. Acetylation at lysine was specified as variable modification, and carbamidomethylation on cysteine was set as fixed modification. Endoprotease ArgC was defined as protease with a maximum of two missed cleavages and the minimum peptide length was set to five. Data was mapped to the ‘El Paco’ protein annotation version 1 and 286 commonly observed contaminants. Initial maximum allowed mass tolerance was set to 4.5 parts per million (ppm) for precursor ions and 20 ppm for fragment ions. Peptide, protein and modification site identifications were reported at a false discovery rate (FDR) of 0.01, estimated by the target/decoy approach ^66^. The label-free algorithm was enabled, as was the “match between runs” option ^67^. A spectrum quality control threshold score >100 and posterior error probability (PEP) <0.01 was defined. The averages of two biological replicates of acetylated-peptide intensities normalized to total H3 and H4 peptide intensities are presented in Fig. 2h,i, and statistical significance was assessed by a two way *student’s t*-test. For fold-change, relative ion intensities of TSA-treated samples were compared to both DMSO and Na-Butyrate.

### Relative *eud-1* expression by Reverse Transcription-quantitative PCR (RT-qPCR)

Worm pellets (25-100 µl) collected at the indicated time points from each culture condition were freeze-thawed 3x between liquid nitrogen and a 37°C heat block, then re-suspended in 500 µl Trizole (Ambion cat. #15596026). RNA was extracted following the manufacturer’s protocol, then purified using a Zymo RNA clean & concentrator-25 columns. RNA was eluted in 50 µl water and quantified by nanodrop (A260/280). 1 µg RNA was used for reverse transcription with SuperScript II (Thermo Fisher cat. #18064071) following manufacturers protocols. Template RNA was degraded by the addition of 40 µl base (150 mM KOH, 20 mM Tris) for 10 minutes, 99°C. cDNA pH was subsequently neutralized by 40 µl acid (150 mM HCl), then diluted by 100 µl TE (200 µl final volume, 1:100 dilution). Four µl cDNA was used as input for 10 µl qPCR reactions with 2x Luna Universal qPCR Master Mix (NEB cat. #M3003X) and 0.25 µM forward and reverse primers (Extended Data Table 1) on a LightCycler 480/II 384 (Roche, serial #6073). Four technical replicates were performed per biological replicate qPCR primer set. Standard quantitative PCR thermocycling conditions were used: 95°C for 5 minutes to denature, followed by 45 cycles of 95°C for 10 seconds (s) (4.8°C/s), 60°C for 10s (2.5°C/s), and 72°C for 10s (4.8°C/s). After qPCR, a thermal melting profile was also obtained for each primer set and verified for whether it was a single peak.

For all experiments quantifying *eud-1* expression by RT-qPCR, threshold Ct values were compared to housekeeping genes *cdc-42* and *y-45* to obtain 2^Δ Ct^, and the geometric mean was calculated for each time point, then normalized to t’0 = eggs, representing 2^-ΔΔCt^, or fold change relative to t’ = 0. Error bars represent S.E.M. from 3-5 biological replicates, and a one-tailed *student’s t*-test between agar and liquid was performed for assessing statistical differences.

### ChIP-qPCR

Nuclear fractionation, chromatin digestion, and immunoprecipitation (IP) were performed as previously described in Werner et al., 2018 with an additional pre-clear step prior to IP with washed, unconjugated beads for 30 minutes. This additional step was empirically determined to yield mildly greater enrichment *vs*. background (Input) compared to the original protocol, or an additional step of hydroxyapatite (hap) nucleosome purification. A 1M salt wash yielded greater signal *vs*. background, however it enriched for multivalent antibody binding, i.e. higher-order nucleosomes, compared to the other conditions. After pre-clear, ∼10 µg of input (fully saturating beads) was used for 10-minute incubations with antibody-conjugated Dynabeads (Thermo Fisher cat. #10004D) 5 µg Ab/20 µl beads, followed by five wash steps and three tube transfers. Coprecipitated DNA was purified by 0.4 mg/ml Proteinase K digestion for two hours, 55°C, and AMPure XP bead purification (cat. #A63881), and resuspended in 50 µl TE buffer.

Quantitative PCR (qPCR) was performed on a 1:100 dilution of co-precipitating DNA with four technical replicates, following the parameters discussed in *eud-1* qPCR (see above), and the average Ct was normalized according to the ‘% Input’ method (Haring et al., 2007) as follows:

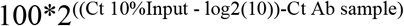

Statistical significance was determined by a one-way *student’s t*-test (based on phenotype, WB and LC-MS/MS results).

### Developmental Delay

*P. pacificus* (PS312) and *C. elegans* (N2) were bleach-synchronized, then eggs were incubated 24 hours in M9 buffer with rotation at room temperature, in order to further synchronize to the L2/J2 stage with starvation. Afterwards, animals were aliquoted into 10 ml standard S-Medium liquid cultures with 66 µM TSA (100 µl) or 100 µl DMSO, and observed at the indicated time points with a 40x oil immersion objective on an upright light microscope (Zeiss Axioscope 5). Stages were identified and scored based on vulva and mouth development, n = 2 (for both nematode species).

*Drosophila* embryos were collected for 3h on standard grape juice agar egg-laying medium, and transferred individually to culture vials. The genotype of all flies used was *w*^*1118*^ (Bloomington Drosophila Stock Center #3605). Flies were cultured on standard medium containing cornmeal, yeast, agar, and molasses, plus 0.1% propionic acid and 0.1% Tegosept for mold inhibition. Additionally, the medium contained either TSA dissolved in DMSO, or DMSOTSA and DMSO-only contro lonly as a control. These were added to the medium at a concentration of 0.34% (v/v) once it had cooled to approximately 50°C. This resulted in a final concentration of 10 µM TSA in the TSA-containing medium. For each condition (10 µM TSA and DMSO-only control), four replicates were established. Each replicate consisted of a vial with 50 embryos each, for a total of 200 embryos per treatment. Beginning 118h after transfer of embryos (118h – 121h post-laying), pupae and adults were counted in each vial every 24h. Flies were maintained at 25°C and 60% humidity on a 16:8 light:dark cycle, and counting occurred 3h after lights on. Pupae were counted through Day 13 (thereafter, any new pupae could conceivably represent progeny of the original flies). Flies were sexed and counted until all replicates of both treatments stopped producing new adults, which occurred on Day 16. Each day, all new adults were frozen at -80°C for use in Western Blots.

Statistical analysis was performed in R. Differences in time to pupation and to eclosion between TSA-treated flies and DMSO-treated controls were analyzed with two-factor ANOVA, accounting for variation between replicates (Model: time ∼ replicate + treatment).

In *Drosophila* raised in media containing 10 µM TSA, the average time to pupation was 8.68 days (95% CI: 8.33 - 9.03) and the average time to eclosion was 12.72 days (95% CI: 12.43 – 13.02). In control flies treated with DMSO only, the average time to pupation was 7.35 days (95% confidence interval: 7.22 – 7.48) and the average time to eclosion was 11.56 days (95% CI: 11.43 – 11.68). Thus, flies exposed to TSA took significantly longer to develop to pupation *(p* < 0.00001, two-way ANOVA) and to eclosion *(p* < 0.00001, two-way ANOVA). The average developmental delay in TSA-treated flies was 1.33 days over the period between laying and pupation and 1.16 days over the entire life cycle from laying to eclosion, indicating that TSA delays development in the larval stages of *Drosophila* but not the pupal stage. We found that flies raised in TSA were significantly less likely to survive from laying to pupation than the DMSO-only controls (56/200 survivors in TSA vs. 142/200 in DMSO,*p* < 0.00001, two-sample chi-square test with Yates correction). However, between pupation and eclosion, the survival rate did not differ significantly between the treatments (43/56 survivors in TSA vs. 124/142 in DMSO, *p* = 0.105, two-sample chi-square test with Yates correction), further suggesting that the main developmental effects of TSA occur prior to pupation. Consistent with previous results (Pile *et al*. 2001) we found that the surviving adults in the TSA-treated group were significantly more likely to be female than DMSO-treated controls (26/43 females in TSA vs. 51/124 in DMSO, *p* = 0.044, two-sample chi-square test with Yates correction).

### Data availability

All nematode strains used in this study are available from the corresponding author Ralf J. Sommer. All other materials (oligonucleotide primers and antibodies) used in this study were purchased from vendors. The ChIP-seq datasets generated during this study are available at the National Center for Biotechnology Information Sequence Read Archive (NCBI SRA) data base under the accession PRJNA628502, and can be accessed with the following link: https://www.ncbi.nlm.nih.gov/sra/PRJNA628502 (released at date of publication). The mass spectrometry proteomics data have been deposited to the ProteomeXchange Consortium via the PRIDE partner repository with the dataset identifier PXD018940 (released at date of publication).

## Acknowledgements

We would like to thank Silke Wahl for guidance on histone LC-MS/MS and loading digested peptides on C18 columns. We would like to acknowledge Hanh Witte and Bogdan Sieriebriennikov for creating and providing the *eud-1* CRISPR mutant. We would also like to thank Talia L. Karasov, James Lightfoot, and Tess Renahan for critical reading of our manuscript, and all members of the Sommer and Werner Laboratories. Funding was generously provided by The Max Planck Society and the School of Biological Sciences at the University of Utah

## Author Contributions

M.S.W. and R.J.S. designed experiments; M.S.W. and T.L. performed reciprocal transplant and RT-qPCR experiments. M.S.W. and T.T. performed GMM. M.S.W. and T.L. performed RT-qPCR. M.S.W. extracted histones and performed nematode WBs, and prepared digested peptides for LC-MS/MS. M.F.W. performed LC-MS/MS with supervision from B.M. T.K. performed all fly experiments. R.J.S. and M.S.W. provided resources. Writing by M.S.W. and R.J.S. with input from all authors.

## References

1. Yan, H. et al. Eusocial insects as emerging models for behavioural epigenetics. Nat Rev Genet 15, 677–688 (2014).

2. Sommer, R. J. et al. The genetics of phenotypic plasticity in nematode feeding structures. Open Biol 7, 160332 (2017).

3. DeWitt, T. J. & Scheiner, S. M. Phenotypic Plasticity: Functional and Conceptual Approaches. (2004).

4. Pigliucci, M. Phenotypic Plasticity: Beyond Nature and Nurture. (Johns Hopkins University Press, 2001).

5. Nijhout, H. F. Development and evolution of adaptive polyphenisms. Evol Dev 5, 9–18 (2003).

6. Müller, G. B. Evo-devo: extending the evolutionary synthesis. Nat Rev Genet 8, 943–949 (2007).

7. West-Eberhard, M. J. Developmental Plasticity and Evolution. (Oxford University Press, 2003).

8. Laland, K. N. et al. The extended evolutionary synthesis: its structure, assumptions and predictions. Proc Royal Soc B Biological Sci 282, 20151019 (2015).

9. Valena, S. & Moczek, A. P. Epigenetic Mechanisms Underlying Developmental Plasticity in Horned Beetles. Genetics Res Int 2012, 576303 (2012).

10. Bento, G., Ogawa, A. & Sommer, R. J. Co-option of the hormone-signalling module dafachronic acid-DAF-12 in nematode evolution. Nature 466, 494–497 (2010).

11. Werner, M. S., ClaaBen, M. H., Renahan, T., Dardiry, M. & Sommer, R. J. Adult Influence on Juvenile Phenotypes by Stage-Specific Pheromone Production. Iscience 10, 123–134 (2018).

12. Lightfoot, J. W. et al. Small peptide-mediated self-recognition prevents cannibalism in predatory nematodes. Science 364, 86–89 (2019).

13. Bose, N. et al. Complex Small-Molecule Architectures Regulate Phenotypic Plasticity in a Nematode. Angew Chem-ger Edit 124, 12606–12611 (2012).

14. Renahan, T. & Sommer, R. J. Nematode Interactions on Beetle Hosts Indicate a Role of Mouth-Form Plasticity in Resource Competition. Frontiers Ecol Evol 9, 752695 (2021).

15. Renahan, T. et al. Nematode biphasic ‘boom and bust’ dynamics are dependent on host bacterial load while linking dauer and mouth-form polyphenisms. Environ Microbiol 23, 51025113 (2021).

16. Werner, M. S. et al. Environmental influence on Pristionchus pacificus mouth form through different culture methods. Sci Rep-uk 7, 7207 (2017).

17. Mather, K. & Winton, D. D. Adaptation and Counter-Adaptation of the Breeding System in Primula: THE NATURE OF BREEDING SYSTEMS. Ann Bot-london 5, 297–311 (1941).

18. Sommer, R. J. Phenotypic Plasticity: From Theory and Genetics to Current and Future Challenges. Genetics 215, 1–13 (2020).

19. Ragsdale, E. J., Müller, M. R., Rodelsperger, C. & Sommer, R. J. A Developmental Switch Coupled to the Evolution of Plasticity Acts through a Sulfatase. Cell 155, 922–933 (2013).

20. Sommer, R. J. Pristionchus pacificus: A Nematode Model for Comparative and Evolutionary Biology. vol. 11 (2015).

21. Rada-Iglesias, A. et al. A unique chromatin signature uncovers early developmental enhancers in humans. Nature 470, 279–283 (2011).

22. Simola, D. F. et al. Epigenetic (re)programming of caste-specific behavior in the ant Camponotus floridanus. Science 351, aac6633 (2016).

23. Ozawa, T. et al. Histone deacetylases control module-specific phenotypic plasticity in beetle weapons. Proc National Acad Sci 113, 15042–15047 (2016).

24. Serobyan, V. et al. Chromatin remodelling and antisense-mediated up-regulation of the developmental switch gene eud-1 control predatory feeding plasticity. Nat Commun 7, 12337 (2016).

25. Yoon, S. & Eom, G. H. HDAC and HDAC Inhibitor: From Cancer to Cardiovascular Diseases. Chonnam Medical J 52, 1–11 (2016).

26. Li, W. & Sun, Z. Mechanism of Action for HDAC Inhibitors—Insights from Omics Approaches. Int J Mol Sci 20, 1616 (2019).

27. Bradner, J. E. et al. Chemical Phylogenetics of Histone Deacetylases. Nat Chem Biol 6, 238243 (2010).

28. Glastad, K. M. et al. Epigenetic Regulator CoREST Controls Social Behavior in Ants. Mol Cell 77, 338-351.e6 (2019).

29. Peleg, S. et al. Life span extension by targeting a link between metabolism and histone acetylation in Drosophila. Embo Rep 17, 455–469 (2016).

30. Nagarajan, S., Benito, E., Fischer, A. & Johnsen, S. A. H4K12ac is regulated by estrogen receptor-alpha and is associated with BRD4 function and inducible transcription. Oncotarget 6, 7305–7317 (2015).

31. Peleg, S. et al. Altered Histone Acetylation Is Associated with Age-Dependent Memory Impairment in Mice. Science 328, 753–756 (2010).

32. Wang, Z. et al. Combinatorial patterns of histone acetylations and methylations in the human genome. Nat Genet 40, 897–903 (2008).

33. Barnes, C. E., English, D. M. & Cowley, S. M. Acetylation & Co: an expanding repertoire of histone acylations regulates chromatin and transcription. Essays Biochem 63, 97–107 (2019).

34. Grzybowski, A. T., Chen, Z. & Ruthenburg, A. J. Calibrating ChIP-Seq with Nucleosomal Internal Standards to Measure Histone Modification Density Genome Wide. Mol Cell 58, 886899 (2015).

35. Sieriebriennikov, B. et al. A Developmental Switch Generating Phenotypic Plasticity Is Part of a Conserved Multi-gene Locus. Cell Reports 23, 2835-2843.e4 (2018).

36. Ambros, V. Control of developmental timing in Caenorhabditis elegans. Curr Opin Genet Dev 10, 428–433 (2000).

37. Serobyan, V., Ragsdale, E. J., Müller, M. R. & Sommer, R. J. Feeding plasticity in the nematode Pristionchus pacificus is influenced by sex and social context and is linked to developmental speed. Evol Dev 15, 161–70 (2013).

38. Howard, R. J. et al. The Ediacaran Origin of Ecdysozoa: Integrating Fossil and Phylogenomic Data. J Geol Soc London jgs2021-107 (2022) doi:10.1144/jgs2021-107.

39. Pile, L. A., Lee, F. W.-H. & Wassarman, D. A. The histone deacetylase inhibitor trichostatin A influences the development of Drosophila melanogaster. Cell Mol Life Sci Cmls 58, 17151718 (2001).

40. Akhtar, A. & Becker, P. B. Activation of transcription through histone H4 acetylation by MOF, an acetyltransferase essential for dosage compensation in Drosophila. Mol Cell 5, 367–75 (2000).

41. Rodrigues, C. P., Shvedunova, M. & Akhtar, A. Epigenetic Regulators as the Gatekeepers of Hematopoiesis. Trends Genet 37, 125–142 (2020).

42. Graff, J. & Tsai, L.-H. Histone acetylation: molecular mnemonics on the chromatin. Nat Rev Neurosci 14, 97–111 (2013).

43. Heintzman, N. D. et al. Histone modifications at human enhancers reflect global cell-type-specific gene expression. Nature 459, 108–112 (2009).

44. Kanno, T. et al. BRD4 assists elongation of both coding and enhancer RNAs by interacting with acetylated histones. Nat Struct Mol Biol 21, 1047–1057 (2014).

45. Kanno, T. et al. Selective Recognition of Acetylated Histones by Bromodomain Proteins Visualized in Living Cells. Mol Cell 13, 33–43 (2004).

46. LeRoy, G., Rickards, B. & Flint, S. J. The Double Bromodomain Proteins Brd2 and Brd3 Couple Histone Acetylation to Transcription. Mol Cell 30, 51–60 (2008).

47. Ediriweera, M. K., Tennekoon, K. H. & Samarakoon, S. R. Emerging role of histone deacetylase inhibitors as anti-breast-cancer agents. DrugDiscov Today 24, 685–702 (2019).

48. Vigushin, D. M. et al. Trichostatin A is a histone deacetylase inhibitor with potent antitumor activity against breast cancer in vivo. Clin Cancer Res Official J Am Assoc Cancer Res 7, 971–6 (2001).

49. Hrgovic, I. et al. The histone deacetylase inhibitor trichostatin a decreases lymphangiogenesis by inducing apoptosis and cell cycle arrest via p21-dependent pathways. Bmc Cancer 16, 763 (2016).

50. Zheng, Y., Thomas, P. M. & Kelleher, N. L. Measurement of acetylation turnover at distinct lysines in human histones identifies long-lived acetylation sites. Nat Commun 4, 2203 (2013).

51. Behera, V. et al. Interrogating Histone Acetylation and BRD4 as Mitotic Bookmarks of Transcription. Cell Reports 27, 400-415.e5 (2019).

52. Samata, M. et al. Intergenerationally Maintained Histone H4 Lysine 16 Acetylation Is Instructive for Future Gene Activation. Cell 182, 127-144.e23 (2020).

53. McNamara, K. J. Heterochrony: the Evolution of Development. EvolEduc Outreach 5, 203218 (2012).

54. Susoy, V., Ragsdale, E. J., Kanzaki, N. & Sommer, R. J. Rapid diversification associated with a macroevolutionary pulse of developmental plasticity. Elife 4, e05463 (2015).

55. Levis, N. A. & Pfennig, D. W. Evaluating ‘Plasticity-First’ Evolution in Nature: Key Criteria and Empirical Approaches. Trends Ecol Evol 31, 563–574 (2016).

56. Stiernagle, T. Maintenance of C. elegans. Wormbook 1–11 (2006) doi:10.1895/wormbook.1.101.1.

57. Goutelle, S. et al. The Hill equation: a review of its capabilities in pharmacological modelling. Fundam Clin Pharm 22, 633–48 (2008).

58. Theska, T., Sieriebriennikov, B., Wighard, S. S., Werner, M. S. & Sommer, R. J. Geometric morphometrics of microscopic animals as exemplified by model nematodes. Nat Protoc 15, 2611–2644 (2020).

59. Werner, M. S. et al. Young genes have distinct gene structure, epigenetic profiles, and transcriptional regulation. Genome Res 28, 1675–1687 (2018).

60. Shechter, D., Dormann, H. L., Allis, C. D. & Hake, S. B. Extraction, purification and analysis of histones. Nat Protoc 2, 1445–1457 (2007).

61. Schindelin, J. et al. Fiji: an open-source platform for biological-image analysis. Nat Methods 9, 676–682 (2012).

62. Rappsilber, J., Mann, M. & Ishihama, Y. Protocol for micro-purification, enrichment, prefractionation and storage of peptides for proteomics using StageTips. Nat Protoc 2, 1896–1906 (2007).

63. Kliza, K. et al. Internally tagged ubiquitin: a tool to identify linear polyubiquitin-modified proteins by mass spectrometry. Nat Methods 14, 504–512 (2017).

64. Cox, J. & Mann, M. MaxQuant enables high peptide identification rates, individualized p.p.b.-range mass accuracies and proteome-wide protein quantification. Nat Biotechnol 26, 1367–1372 (2008).

65. Cox, J. et al. Andromeda: A Peptide Search Engine Integrated into the MaxQuant Environment. JProteome Res 10, 1794–1805 (2011).

66. Elias, J. E. & Gygi, S. P. Target-decoy search strategy for increased confidence in large-scale protein identifications by mass spectrometry. Nat Methods 4, 207–214 (2007).

67. Luber, C. A. et al. Quantitative Proteomics Reveals Subset-Specific Viral Recognition in Dendritic Cells. Immunity 32, 279–289 (2010).

