## extended data figures for "Histone 4 lysine 5/12 acetylation provides a plasticity code with epigenetic memory of environmental exposure"

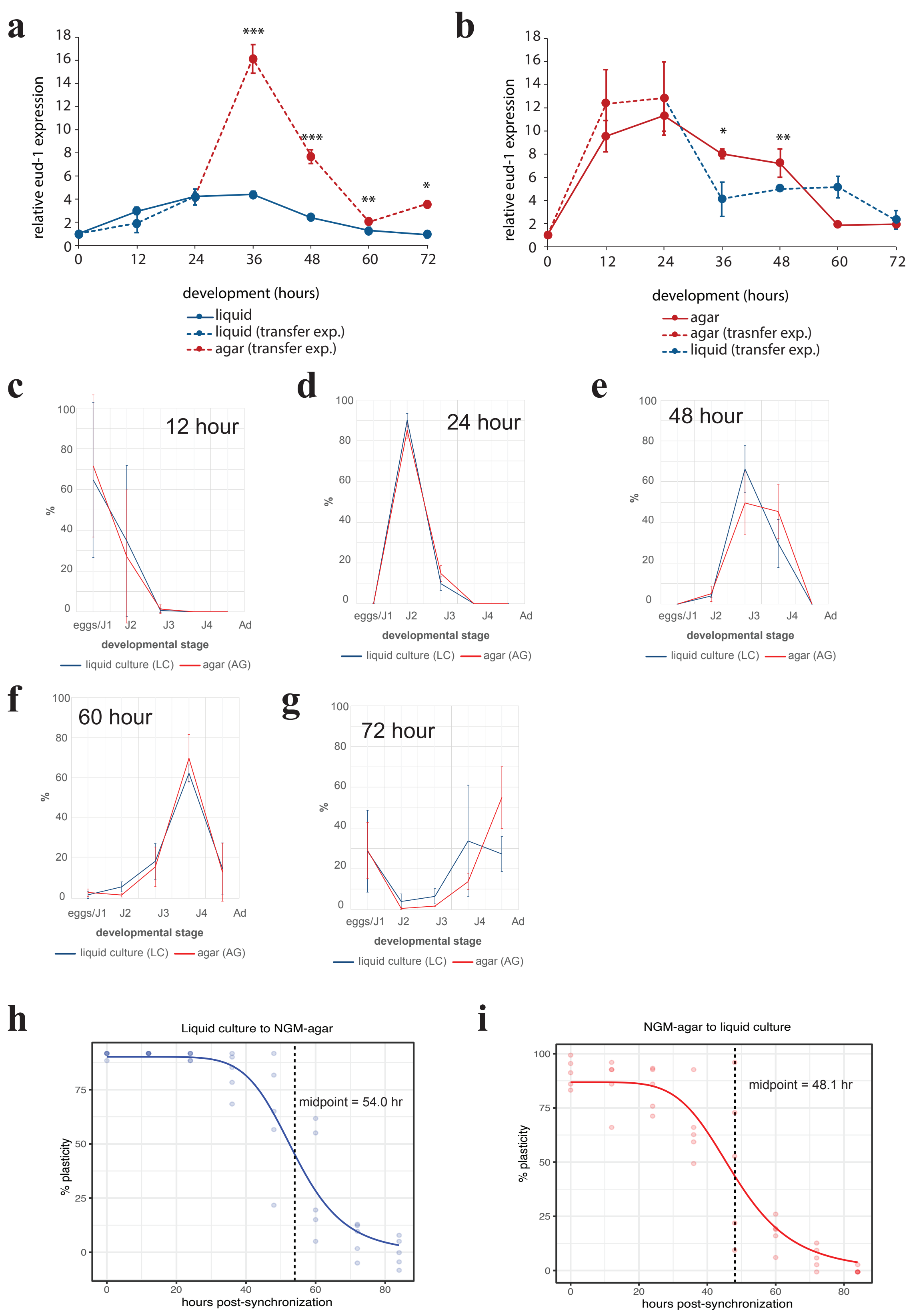


**Extended Data Fig. 1 | Switch gene transcription corresponds to environemntal sensitivity.**
**a,** Transcription of *eud-1* when worms are transferred from liquid culture to NGM-agar, or **b,** in the reverse direction. Relative *eud-1* expression was measured by RT-qPCR (geometric mean of $2^{\Delta Ct}$ relative to *Ppa-cdc-42* and *Ppa-y-45*, normalized to t'=0). Error bars represent S.E.M. for 3-5 biological replicates. Statistical significance was determined for every time point in agar against the equivalent time piont in liquid culture by a one-sided student's t-test, *=p<0.05, **=p<0.01, ***=p<0.001. **c-g,** Developmental stages in agar (red) and liquid culture (blue) after hypochlorite ('bleach') synchronization, n=3, error bars reflect standard deviation. **h,** logistic fit of reciprocal transplant experiments (Fig. 1c) from liquid to NGM-agar and **i,** vice-versa. y-axis (% plasticity) represents %Eu values normalized to the ground-state mouth-form ratios in either environment (see methods for more details).

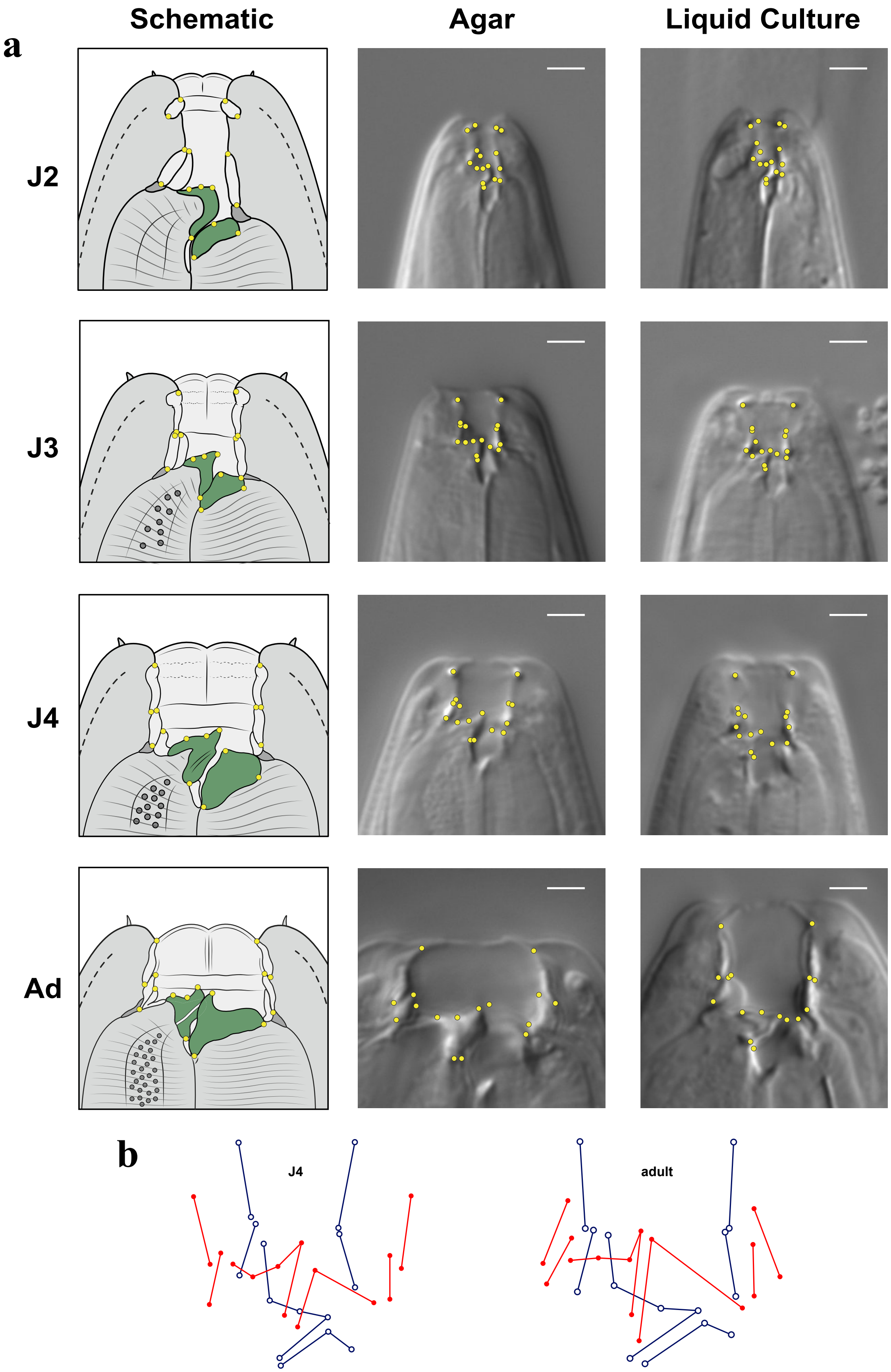


**Extended Data Fig. 2 | Homologous landmarks for quantitative geometric morphometrics**
**a** Homologous structural landmarks used for GM in both NGM-agar and liquid culture at each developmental stage (except for J1). Note, only one morph (Eu) is shown for the adult illustration, and some landmarks in the microscopy images are out of the current focal plane. Scale bar = 5 µm. Schematics drawn in Adobe illustrator, and images taken using 100x 1.4 oil immersion objective with DIC prism on a Zeiss Axio Imager. **b** Wire-frame plots of each morph (red=Eu, blue=St) in J4 and adults. Produced in R according to Theska et al. 2020.

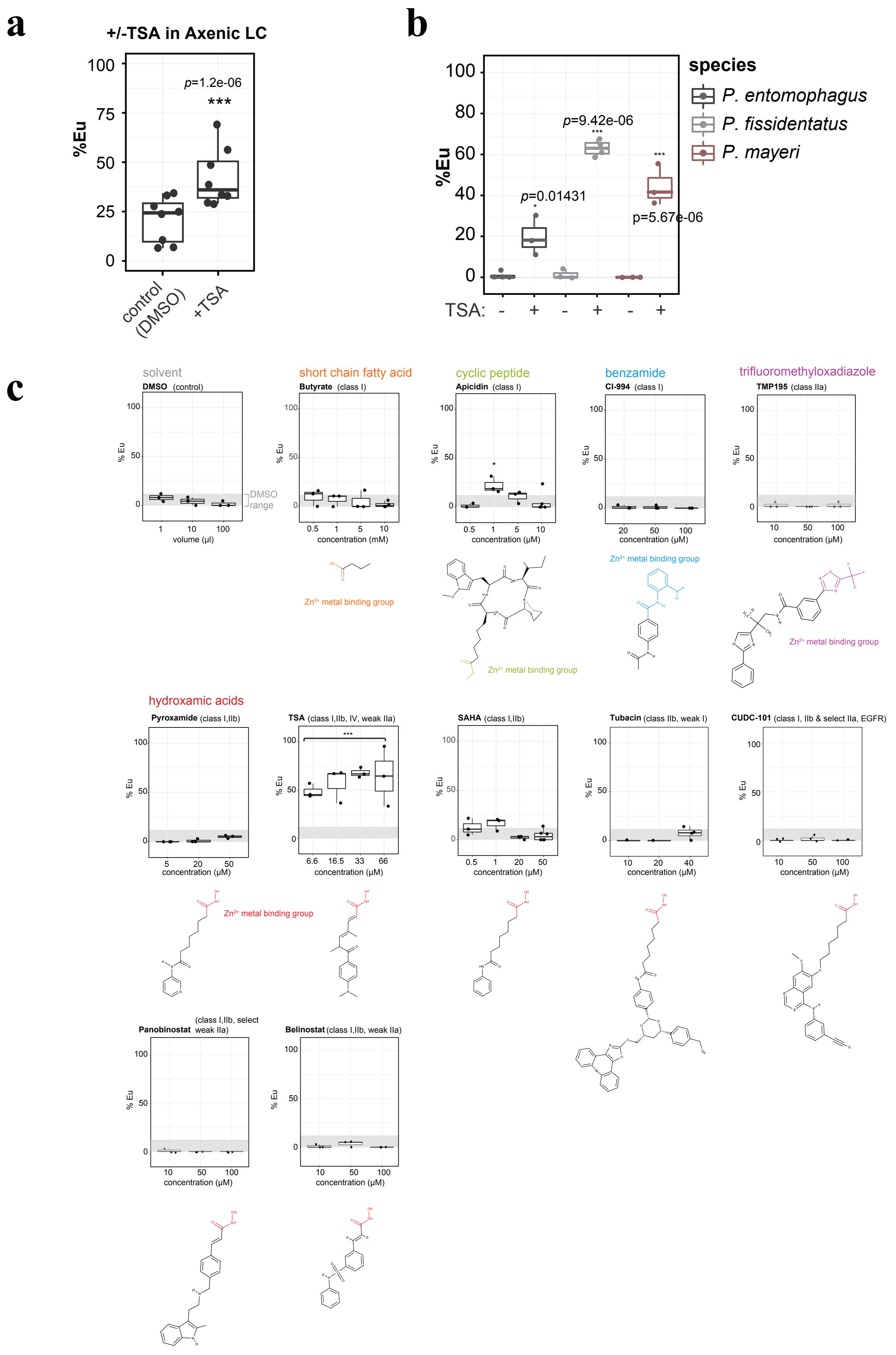


**Extended Data Fig. 3 | TSA has a conserved and specific effect on mouth form. a,** Phenotype of *P. pacificus* in axenic liquid culture +/- TSA. **b,** Different *Pristionchus* species in LC +/- TSA. **c,** Phenotype of *P. pacificus* animals grown in liquid culture with the indicated amounts of HDAC inhibitors. Statistical significance calculated by binomial logistic regression. Samples without TSA were treated with 100 µl DMSO.

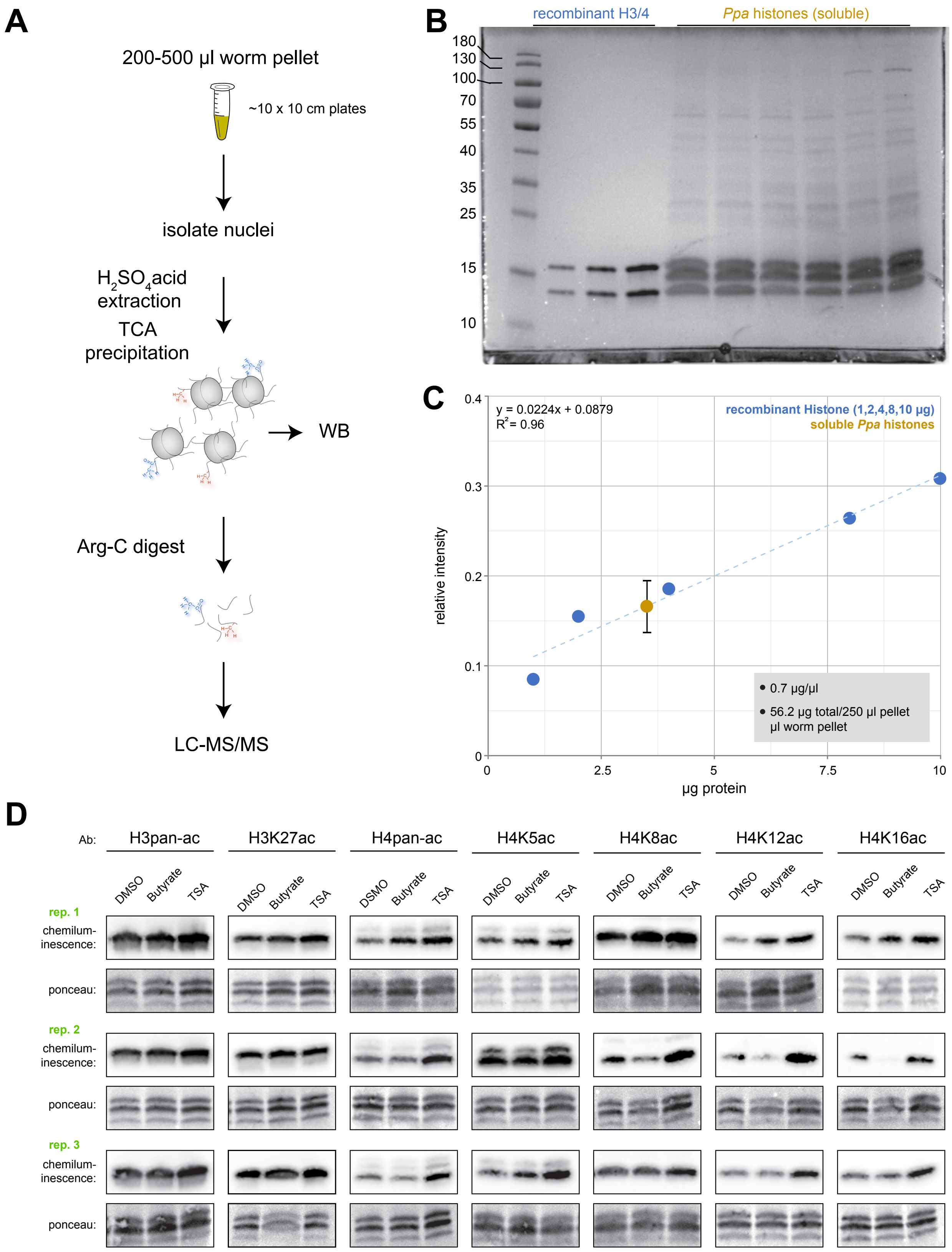


**Extended Data Fig. 4 | Histone Purification and Raw Data for Western Blots.** **a,** Schematic of histone acid-extraction for Western Blot and LC-MS/MS after digestion with Arg-C protease. **b,** Example SDS-PAGE of histone extraction and a recombinant H3/4 calibration curve. Proteins were visualized with Coomasie Brilliant Blue R-250. **c,** Example calibration curve with recombinant histone used to calculate *P. pacificus* histone quantity for WB and LC-MS/MS. **d,** Raw data of chemiluminescent signal from three independent biological replicate WBs for each antibody, and total histone amount transferred to nitrocellulose membranes visualized with Poncaeu Red.

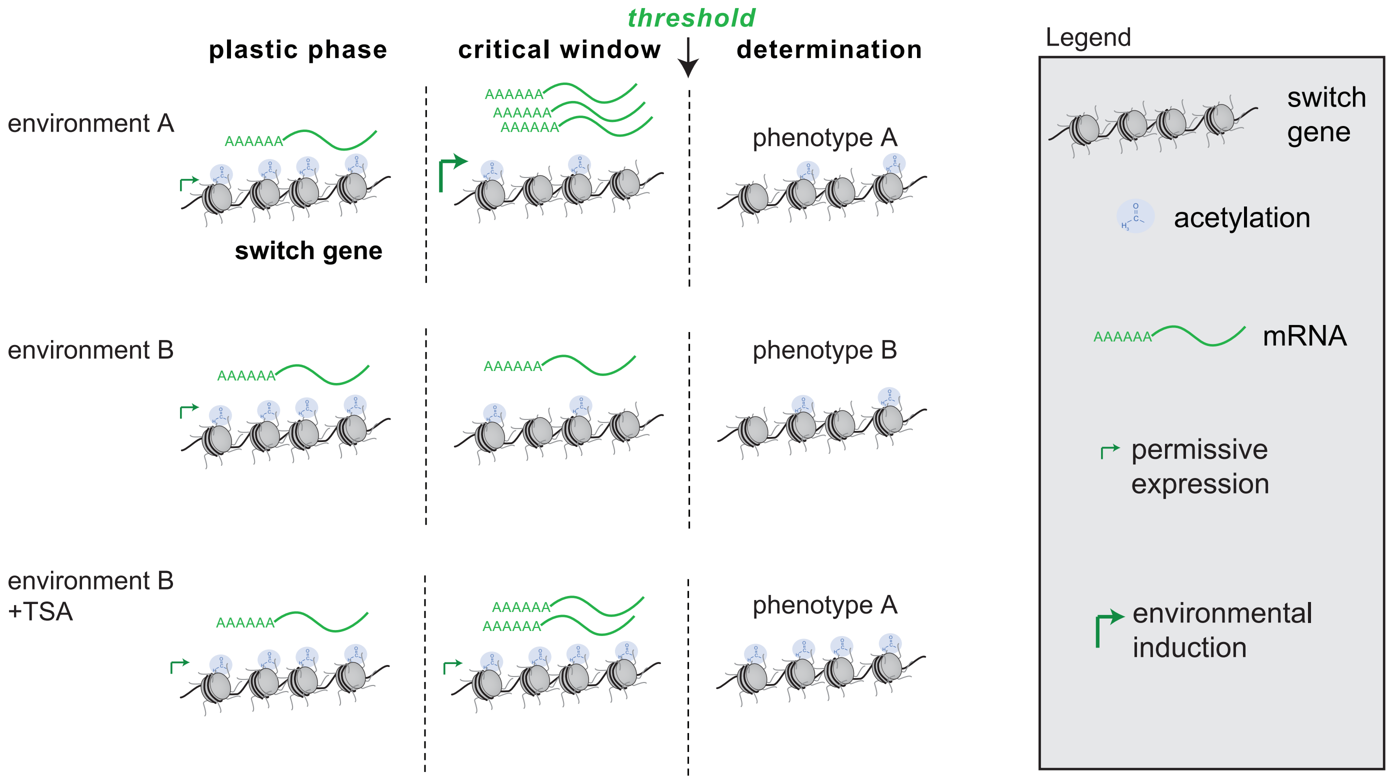


**Extended Data Fig. 5 | Model for establishment of plasticity during the critical window.** Plasticity is established by environment-independent H4K5/12ac and permissive transcription during the critical window. Ultimately H4K5/12 is deacetylated leading toward repression of switch genes and the end of the critcal window. When TSA is added in liquid culture, the fixed juvenile chromatin state of H4K5/12ac prevents switch gene repression.
